# Amplification-free detection of viral RNA by super resolution imaging-based CRISPR/Cas13a System

**DOI:** 10.1101/2021.07.17.452803

**Authors:** Qian He, Qun Chen, Fang Li, Xi Yuan, Chuhui Wang, Changyue Liu, Lidan Xu, Xiaoyun Zhong, Jiazhang Wei, Vijay Pandey, Dongmei Yu, Yuhan Dong, Yongbing Zhang, Lin Deng, Ke Du, Peiwu Qin

## Abstract

RNA detection is crucial for biological research and clinical diagnosis. The current methods include both direct and amplification-based RNA detection. These methods require complicated procedures, suffering from low sensitivity, slow turnaround, and amplification bias. The CRISPR/Cas13a system is a direct RNA detection method via target RNA induced collateral cleavage activity. However, to detect low concentration RNA with CRISPR/Cas13a, target amplification is always required. Herein, we optimize the components of the CRISPR/Cas13a assay to enhance the sensitivity of viral RNA detection which improve the detection limit from 1 pM up to 100 fM. In addition, the integration of CRISPR/Cas13a biosensing and single molecule super resolution imaging is a novel strategy for direct and amplification-free RNA detection. After surface modification, fluorescent RNA reporters are immobilized on the glass coverslip surface and fluorescent signals are captured by total internal reflection fluorescence microscopy (TIRFM), shifting the measurement from spectroscopy to imaging. We quantify the fluorescence signal intensity before and after collateral cleavage of the CRISPR system when viral RNA is present and achieve a detection limit of 10 fM. Therefore, we provide a novel TIRFM-based system to visualize the CRISPR trans-cleavage for direct and robust RNA detection.

## Introduction

RNAs are essential molecules in living cells and the genetic material for certain pathological viruses. The visualization and detection of RNA are important for the understanding of life and clinical diagnosis^1,2^. Various methods have been developed for cellular and viral RNA detection and imaging. Northern blot (known as RNA gel blot)^3,4^ and fluorescence *in situ* hybridization^5,6^ are classical methods to directly quantify and visualize RNA. However, they are labor-intensive and time-consuming with poor sensitivity. Advanced strategies such as RNA microarray^7,8^ and sequencing^8,9^ are superior in terms of fidelity and throughput. The intricate sample pretreatment, complex procedure, and large sample requirement limit the application of these methods. Nucleic acid amplification-based technology (NAAT) exhibits a robust capability for RNA detection, such as real-time polymerase chain reaction (RT-PCR)^10–12^, recombinase polymerase amplification (RPA), loop-mediated isothermal amplification (LAMP) and so on^13–17^. These strategies elevate the sensitivity of RNA detection and have become the gold standard for diagnosing virus-associated diseases^18,19^. NAAT requires multiple procedures, including RNA extraction, reverse transcription, and template replication, which cause RNA loss and false-positive errors, leading to a failure for rapid, accurate, and sensitive detection of contagious RNA virus, such as severe acute respiratory syndrome coronavirus 2 (SARS-CoV-2). Therefore, it is imperative to develop a robust detection method without complicated target amplification.

Clustered Regularly Interspaced Short Palindromic Repeats (CRISPR)/CRISPR-associated protein (Cas) is a bacterial adaptive immune system. Cas13a is an RNA-guided RNA editing enzyme from type VI, class 2 CRISPR/Cas family^20^. CRISPR/Cas13a complex contains Cas13a protein and CRISPR RNA (crRNA), which possesses a target-triggered collateral cleavage after specific crRNA-target RNA recognition^21,22^. This collateral enzymatic activity generates 10^4^ turnovers of fluorescent labeled RNA reporter as a signal amplification mechanism for direct RNA detection^23–25^. However, for molecular diagnosis, CRISPR system is always coupled with front-end target amplification methods to enhance the sensitivity. Efforts have been made towards an amplification-free CRISPR/Cas13a detection system by optimizing crRNA design, usage of multiple crRNAs, or coupling a microfluidic droplet chip with fluorescence microscopy to increase the detection sensitivity^26,27^. However, the combination of super resolution imaging with CRISPR/Cas13a assay for direct RNA detection has not been realized.

In this work, we optimize the CRISPR/Cas13a system and combine the CRISPR/Cas13a assay with total reflection internal fluorescence microscopy (TIRFM) to improve the RNA detection sensitivity for fluorescent reporters. TIRFM is a sensitive fluorescent imaging technique with a single fluorophore capturing capability^28^. The evanescent wave of TIRFM illumination excites the fluorophores on a thin focal plane (~300 nm), reducing the background and providing a high signal to noise ratio (SNR). We immobilize fluorescent RNA reporters onto a coverslip surface with polyethylene glycol (PEG) coating, and the changes of fluorescence intensity before and after Cas13a collateral cleavage is imaged and quantified by TIRFM (**Fig. 1**). With this sensitive amplification-free technique, we can detect 10 fM viral RNA. Finally, we apply this TIRFM-based CRISPR/Cas13a system to the virus mimic throat swab sample with high sensitivity and confirm the feasibility of this method. Thus, we demonstrate a novel amplification-free platform to directly detect RNA without relying on complicated target amplification.

**Fig. 1.**
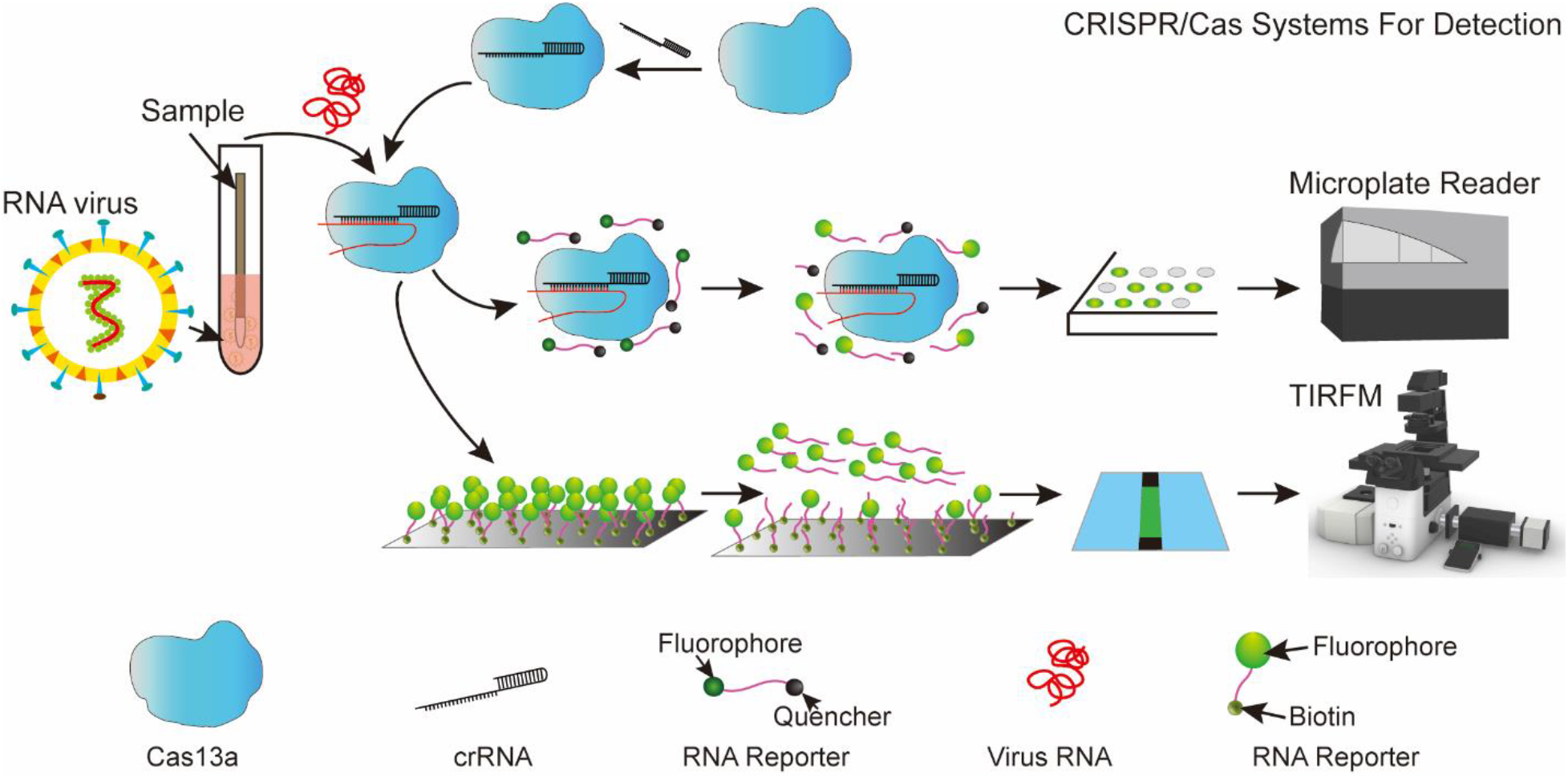
The schematic of viral RNA detection by CRISPR/Cas13a system. We apply CRISPR/Cas13a system to detect the viral RNA directly without amplification. CRISPR/Cas13a system utilizes the bound crRNA to specifically target the viral RNA by base pairing. The Cas13a-crRNA-target RNA complex collaterally cleaves the RNA reporters. The changes of fluorescence signal intensity are detected by microplate reader for off-coverslip assay and TIRFM for on-coverslip assay.

## Results

### CRISPR/Cas13a assay optimization

The CRISPR/Cas13a system consists of multiple components including fluorescent reporters. The catalytic efficiency of 2U is lower than 5U reporters^29^. However, how the length and structure of reporters affect the sensitivity of RNA detection has not been systematically investigated. We vary the RNA reporter length from 5U to 22U to compare the CRISPR/Cas13a assay sensitivity at fixed 10 nM target RNA concentration. All the reporters have the same pair of quencher and fluorophore except the number of inserted uracil nucleotides between them **(supplementary table 1)**. The assay with 22U reporter exhibits the maximal increment of fluorescence intensity (**Fig. 2a**) and achieves a limit of detection (LoD) of 1 pM (**Fig. 2b,2c**), which is ~10 times more sensitive than the 5U case (**supplementary Fig.1**). These results demonstrate that the trans-cleavage activity and detection sensitivity can be enhanced by changing the reporter length. Besides, we find that the fluorescent signal increment remains similar for reporters with various structures (**supplementary Figure 2**). For other components in CRISPR/Cas13a assay, Mg^2+^ concentration within certain range 4 mM to 18 mM barely influences the signal intensity of CRISPR/Cas13a assay when target RNA concentration is between 100 pM and 100 fM, and the LoD is unchanged for the pH between 6.3 to 8.1 (**supplementary Figure 3**).

**Fig. 2.**
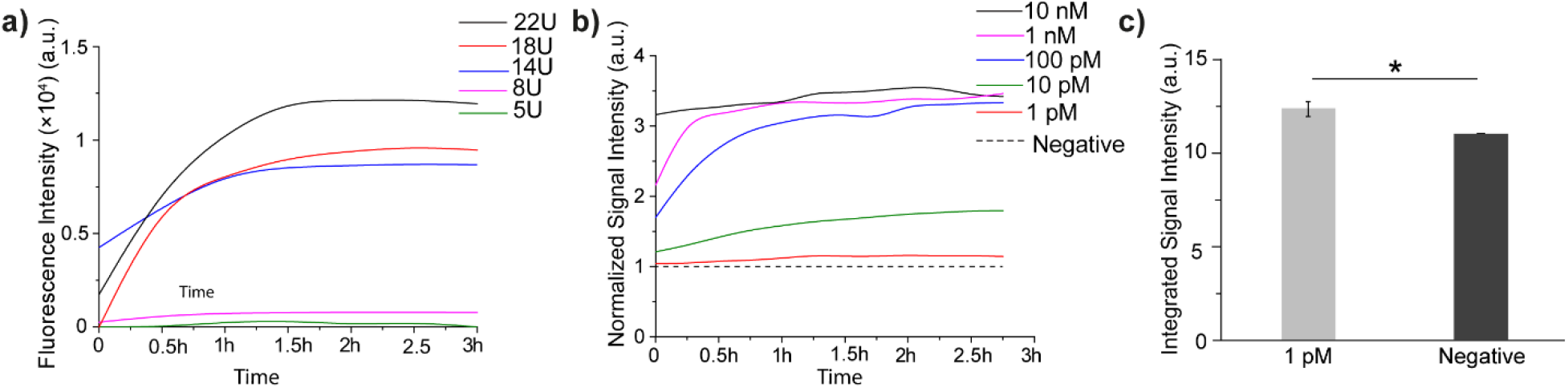
The optimization of RNA reporter. **a)** The increment in signal intensity for CRISPR/Cas13a assay with different RNA reporters. **b)** The signal intensity of the CRISPR/Cas13a assays with 1 pM to 10 nM target RNA using the 22U reporter. **c)** Comparison of integrated signal intensity of CRISPR/Cas13a assay with 1 pM target RNA and negative control. *Student*’s t-test is performed, * indicates *p <* 0.5%.

### TIRFM-based CRISPR/Cas13a system for on-coverslip detection

After optimizing the reporter probe length, we establish a TIRFM-based CRISPR/Cas13a system for RNA sensing. The slide surface is coated with PEG-Biotin and streptavidin to bridge biotin labeled RNA reporter (**Figure. 3a**). The fluorescent signals of all groups are reduced after treatment from images (**Fig. 3b**). We quantify and compare the signal intensities of TIRF images before and after the CRISPR/Cas13a system introduction, the integrated signal intensity of the positive groups exhibits a statistically significant difference after CRISPR/Cas13a introduction (**Fig. 3c**). Higher target RNA concentrations trigger more vigorous trans-cleavage activity, which inversely correlates with the concentration of the target RNA. We use the average fluorescence signal intensity per ROI area to quantify the enzymatic. Higher target RNA concentrations triggers more vigorous trans-cleavage activity, which inversely correlates with the concentration of the target RNA. cleavage of CRISPR/Cas13a since the average intensity per fluorescent puncta between negative and positive groups are similar (**Supplementary Figure 4**).

**Fig. 3.**
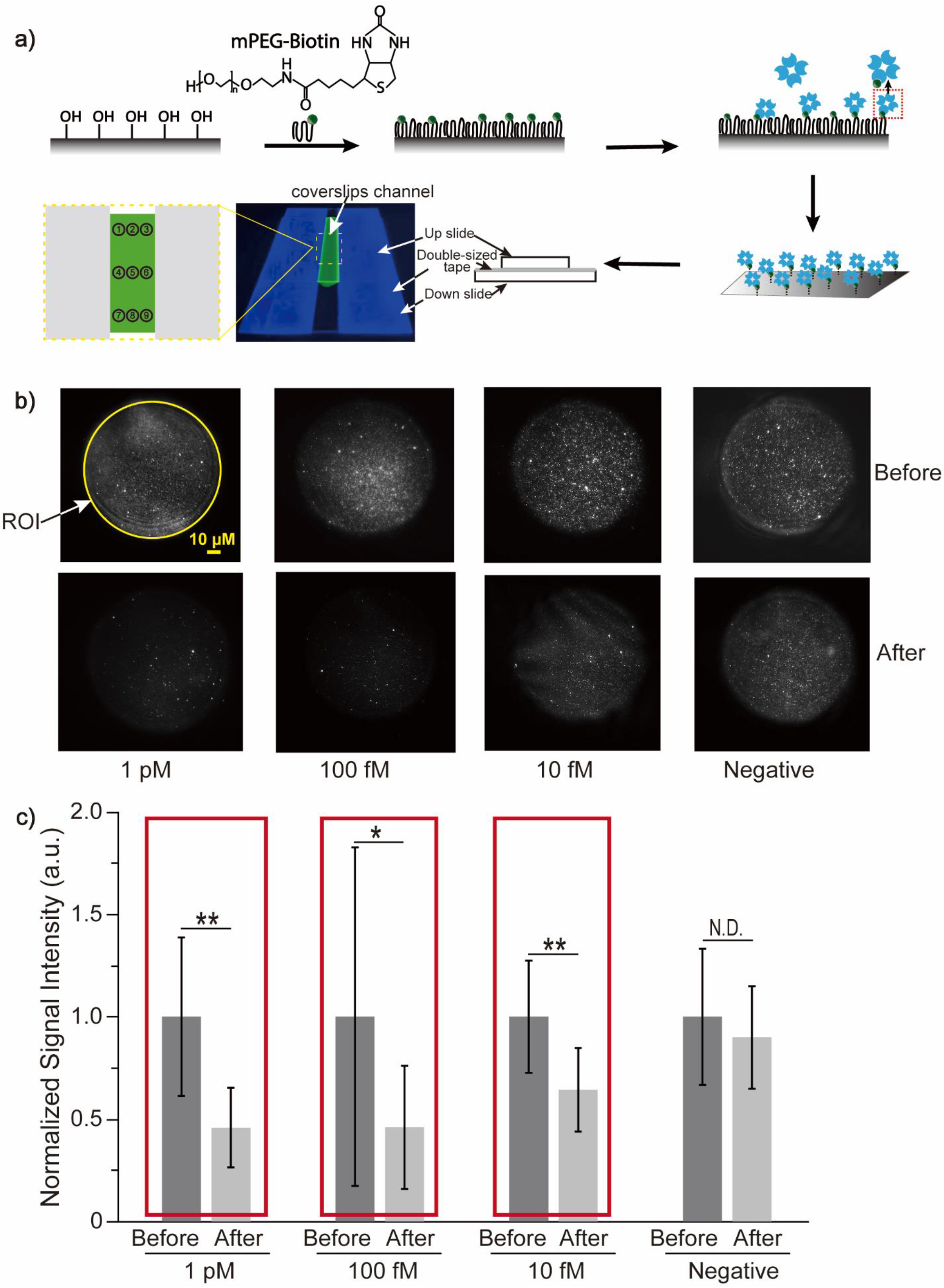
On-coverslip assay for viral RNA detection. **a)** Modification of coverslip for TIRFM-based CRISPR/Cas13a system. The coverslips are cleaned and eroded; next, the surface of coverslip is incubated with mPEG-biotin, and then the fluorescent RNA reporters are immobilized onto a coverslip surface; the channel for on-coverslip assay is created by two coverslips with modified surface facing to each other and are glued by double-sided tape (side view diagram is on the right) and the fluorescent images from nine areas of a channel (yellow dashed line) are captured by TIRFM. **b)** The TIRF images of immobilized fluorescent reporters before and after CRISPR/Cas13a introduction with or without target RNA. The fluorescent signal decreases in a target RNA concentration-dependent manner. ROI: region of interest. **c)** The signal intensity changes are proportional to the target RNA concentration. The LoD of 10 fM is achieved by the combination of CRISPR/Cas13a and TIRFM. *Student*’s t-test is performed, * indicates *p <* 0.05, ** indicates *p <* 0.01, N.D. indicates no difference.

### TIRFM-based CRISPR/Cas13a system for positive mimic sample

To confirm the practicability of the TIRFM-based CRISPR/Cas13a system, we create a SARS-CoV-2 positive mimic sample. The mimic positive sample is *in vitro* transcribed RNA from the SARS-CoV-2 genome plasmid mixed with the throat swab lysate from a healthy individual. The transcribed RNA has been quantified by RT-PCR with a cycle threshold (Ct) value in the range of 23 to 25. The mixture is directly used for CRISPR/Cas13a assay without RNA purification and amplification. The TIRF images show that the fluorescent signal is decreased for both positive mimic and negative control after CRISPR/Cas13a system introduction (**Fig. 4a**). The average fluorescence signal intensity of the positive group shows a significant reduction before and after treatment, which is not observed in the negative control (**Fig. 4b**).

**Fig. 4.**
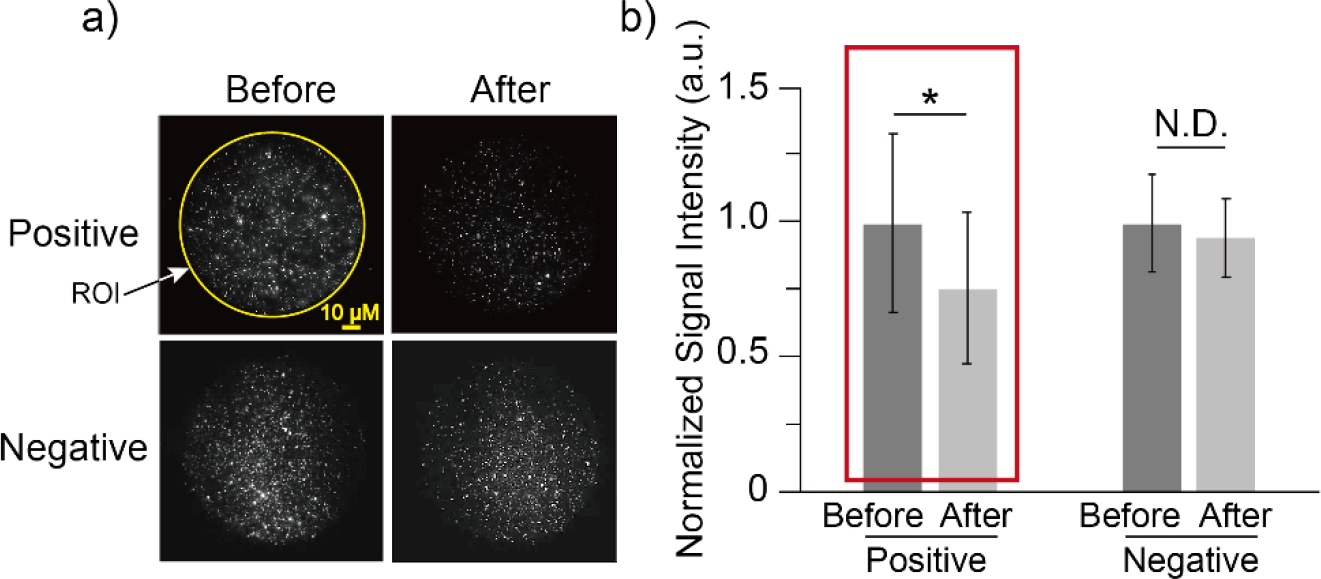
On-coverslip assay for SARS-CoV-2 positive mimic sample. **a)** The TIRF images of the positive mimic and negative control. Reduced fluorescent signal is observed after CRISPR/Cas13a introduction. **b)** The significant difference of signal intensity is observed for the SARS-CoV-2 positive mimic sample. *: *p <* 0.05. N.D. indicates no difference.

## Discussion

Cas13a shows a preference for polyU among the homopolymers such as PolyA, PolyU, PolyG and PolyC and di-nucleotide reporters such as rCrU, rGrU, rArU and rUrU^30^. The catalytic efficiency of 2U is much lower than using 5U as the trans-cleavage substrate of LbuCas13a^31^. Our results show that the 22U reporter exhibits the maximum increment of fluorescent intensity after background subtraction compared with the 5U reporters. This improvement increases the LoD by 10 folds. Reporter longer than 22U will reduce the quenching efficiency and increase the background fluorescence intensity.

Considering the TIRFM has high SNR in fluorescence imaging, and it has been extensively used to study macromolecular interaction ^32,33^, real-time visualization of DNA replication^34^, and identification of single-nucleotide polymorphism^35,36^. We combine the CRISPR/Cas13a system with TIRFM to detect the viral RNA and achieve the LoD of 10 fM without amplification. Our sensitivity is higher than the conventional liquid phase CRISPR/Cas13a assay^25^. The median Ct values of qPCR for asymptomatic, pre-symptomatic, typical symptomatic, and atypical symptomatic residents, were 25.5, 23.1, 24.2, and 24.8, respectively^37^. We detect the mimic positive sample with the comparable RNA concentration as clinical specimens and the Ct value is in the range of 23 to 25 used for our TIRFM-based CRISPR/Cas13a system, which partially covers the range of the viral loading in clinical scenario. This amplification-free platform might be helpful for direct pathogen detection and infection control. Furthermore, to detect lower pathogen loading sample, we could improve the detection sensitivity by introducing brighter reporters such as quantum dots or isothermal DNA amplification for on-coverslip CRISPR/Cas13a assay^38,39^. In addition, artificial intelligence can be introduced into our detection platform for binary classification based on TIRF images. TIRFM-based microarray can be interfaced with CCD cameras or other read-out devices to develop portable devices for rapid RNA detection without amplification^40,41^. Our novel TIRFM-based CRISPR/Cas13a RNA detection platform improves the LoD to 10 fM, which can play a critical role in containing pandemic threat in the future.

## Methods

### Target RNA and crRNA

We pick the 1900 nt sequence from the 5’ terminal of the S gene as the target RNA for CRISPR/Cas13a detection (**Supplementary Table 2**). S gene encodes spike glycoprotein and contains the conserved unique sequences of SARS-CoV-2 compared to other coronaviruses. The target fragment, with T7 promotor (TAATACGACTCACTATAGGGAGA) and polyA tail (AAAAAAA) at the 5’ and 3’ terminal respectively, is inserted into plasmid pUC57. crRNAs are designed according to the 1900 nt target RNA sequence. In addition, we design a shorter 99 nt target from the1900 nt fragment, which has been used previously^42^ (**Supplementary Table 2**). The plasmid and 99 nt RNA target from the S gene are purchased from GeneScript, China. The crRNAs are purchased from Sangon Biotech, China (**Supplementary Table 3**).

### Target fragments amplification and *in vitro*transcription

The target fragments are amplified by PCR using primers (Sangon Biotech, Shanghai, China) and PCR amplification kit (Takara Bio, Japan). The PCR amplifies the DNA fragments including the 1900 bp target, T7 promotor, and polyA tail. The primer sequences are: Forward 5’-CAGATCTTAATACGACTCACTATAGGGAG −3’; Reverse 5’-GGATCCTTTTTTTGCCAA\ GTAGGAG −3’. The PCR is run on a series multi-Block thermal Cycler PCR instrument (LongGene, China) with the following procedures: 95 °C for 5 min (1 cycle); 95 °C for 30 s, 59 °C for 45 s, and 72 °C for 2 min (35 cycles); 72 °C for 5 min (1 cycle); and 4 °C for storage. Agarose gel electrophoresis is used to confirm the DNA size, and a quick gel extraction kit (PureLink, invitrogen, USA) is used to purify the amplified DNA fragments (**Supplementary Figure 5**). *In vitro* transcription is conducted using T7 RNA polymerase kit (D7069, Beyotime, China), NTP (ATP, GTP, CTP, GTP from Thermo Scientific, USA), and the purified DNA fragments as the template. The denaturing formaldehyde gels are used to verify the size of the target RNA (**Supplementary Figure 5**).

### ***Lwa***Cas13a protein purification

PC013 plasmid of *Leptotrichia wadeii* Cas13a (#90097, Addgene, USA) is transformed into Rosetta (DE3) pLysS competent cells for protein expression. The expression and purification procedures follow the published protocol^43^. In brief, 0.5 mM IPTG is added to induce the *Lwa*Cas13a expression at an OD_600_ of 0.6. The cells are cooled down to 18 °C and incubated 16 h for protein expression. Subsequently, the cells are collected by centrifugation. Lysis buffer (20 mM Tris-HCl, 500 mM NaCl, 1 mM DTT, pH 8.0) with 1 tablet of protease inhibitors, lysozyme, and benzonase nuclease is applied to rupture the centrifugated cell by sonication. The lysate is cleared by centrifugation, and the supernatant is filtrated and applied to HisTrap™ FF (17-5319-01, GE healthcare, USA) for preliminary purification. Further purification is performed through cation exchange (Mono S, GE Healthcare Life Sciences, USA), and the molecular weight of *Lwa*Cas13a protein (**Supplementary Figure 6**) is verified by SDS-PAGE gel. Proteins are flash-frozen by liquid nitrogen and stored at −80 °C.

### Off-coverslip detection assay and optimization

The CRISPR/Cas13a assay includes 150 μl of 20 nM purified *Lwa*Cas13a, 10 nM crRNA, 167 nM RNA reporters with quencher and fluorophore on two terminals (GeneScript, China), 2 unit of murine RNase inhibitor (New England Biolabs), varying amounts of target RNA, and reaction buffer (40 mM Tris-HCl, 60 mM NaCl, 4 mM MgCl_2_, pH 7.3). Reactions proceed for 5–20 min at room temperature followed by fluorescent measurement with a microplate reader (SPARK, TECAN, Switzerland). In order to improve the LoD, we optimize the assay components including the length of RNA reporters, the pH, the magnesium ion concentration, and the number of crRNAs (**Supplementary Table 3**). A 96 well-plate is used for off-coverslip detection. The fluorescence signal intensity of each well is measured every 15 minutes after 3 seconds of shaking. Excitation and emission wavelengths are 485 nm and 535 nm, respectively. The parameters are the average of 10 flashes for each point, and gain is 55. All data are normalized by negative control, then integrated for two sample T-Test analysis.

### Coverslip cleaning and coating

The surface of the glass coverslip (No.1.5, 24 × 30 mm and 24 × 40 mm, VWR, Germany) is modified and manipulated according to the previous protocol^44^. Glasses are cleaned and eroded by acetone for 30 min and 3 M KOH for 1 h, respectively. After rinsing with distilled water, the glasses are dried by nitrogen gas and immersed into the aminosilane solution with 100 mL methanol, 5 mL glacial acetic acid, 1 mL N-(2-Aminoethyl)-3-Aminopropylmethyldimethoxysilane (BIOFOUNT, China) for 10 min in the dark followed by 1-2 min sonication, then 10 min in the dark again. After rinsing and drying, 4 mg of mPEG-biotin (MW 5000, aladdin, China) and 115 mg mPEG-Succinimidyl Valerate (MW 5000, Seebio, China) in 490 μl sodium bicarbonate buffer is added on the surface of glasses for overnight incubation. Finally, the glasses are put into the container and stored at −80 °C after rinsing and drying.

The coverslip channel is created by placing the smaller coverslip (No.1.5, 24 × 30 mm) on the top of the larger one (No.1.5, 24 × 40 mm) with the modified surface facing to each other and connected by two double-sided strips. 1 mg/mL streptavidin (Merck, Germany) is pipetted into the channel for 10 min incubation. The residual reporter is flushed away. Then, the channel is incubated with 10 μM 22U RNA reporters (Supplementary Table 1) and rinsed with water.

### TIRFM imaging

The fluorescent signal is imaged by TIRFM (Ti2, Nikon, Japan) with 80° incident angle and 500 ms of exposure time. The laser power is 1 mW and the gamma parameter is 1, and the LUT intensity range is from 1 to 2000. sCMOS has 1024 × 1024 pixels. More than 10 images from different areas of the individual channel are taken. For the positive group, the CRISPR/Cas13a system with varying amounts of target RNA is flowed into the channel for 5 min incubation to cleave the immobilized RNA reporters on the coverslip surface. The negative group is CRISPR/Cas13a system without target RNA or RNA from other species. We draw the same size of ROI for each TIRF image and measure the average signal intensity of particle in ImageJ. Therefore, we measure the average fluorescence signal intensity of all ROIs. All data are normalized by negative control, then integrated for two sample T-Test.

### SARS-CoV-2 RNA of mimic clinical sample

The clinical sample is mimicked by mixing the *in vitro* transcribed RNA of SARS-CoV-2 and the throat swab lysate from a healthy person. The throat swab is stored using a single-use virus sampling tube with preserving fluid (Biocomma, Shenzhen, China) and inactivated at 56°C for 30 min. RNA is released by pathogen inactivated, nucleic acid extraction-free, direct-to-PCR buffer with proteinase K (Ebio, Shenzhen, China) and mixed with *in vitro* transcribed viral RNA. The lysate is added directly to the CRISPR/Cas13a system for detection without extraction and purification. Preserving fluid and releasing buffer without viral RNA are added to CRISPR/Cas13a system as the negative control.

## Author contributions

P.Q. supervised the study. Q.H. and Q.C. performed the experiment, collected data, analyzed data and wrote the first draft of the manuscript. F.L., X.Y., L.X., and X.Z. purified the protein. J.W., V.P., D.Y., Y.D., Y.Z., L.D. and K.D. helped revise the manuscript.

