## Supplementary Information for "Amplification-free detection of viral RNA by super resolution imaging-based CRISPR/Cas13a System"

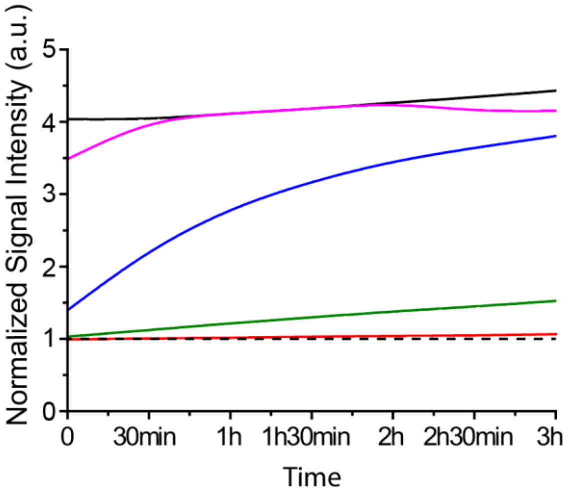


**Supplementary Figure 1.** Target RNA from 1 pM to 10 nM is detected using 5U reporter.

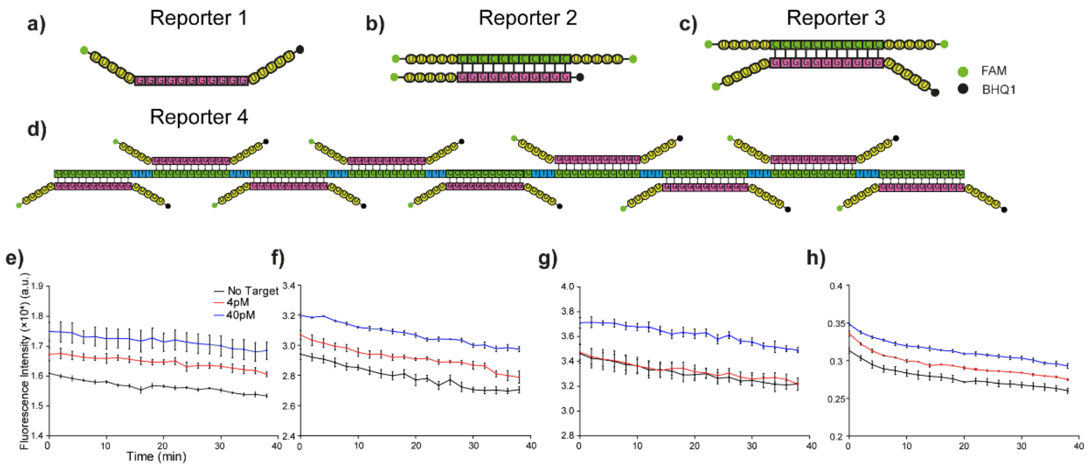


**Supplementary Figure 2. a)** Reporter 1 - BHQ1-5U-11G-5U-FAM. **b)** Reporter 2 - FAM-5U-11C-5U-FAM paired with BHQ1-5U-11G-FAM. **c)** Reporter 3 - FAM-5U-11C-5U-FAM paired with BHQ1-5U-11G-5U-FAM. **d)** Reporter 4 - nine 11C-3T repeats paired with BHQ1-5U-11G-U-FAM. **e - h)** The signal intensity of the CRISPR/Cas13a assays with 4 pM and 40 pM target RNA using the reporter 1 to 4 respectively.

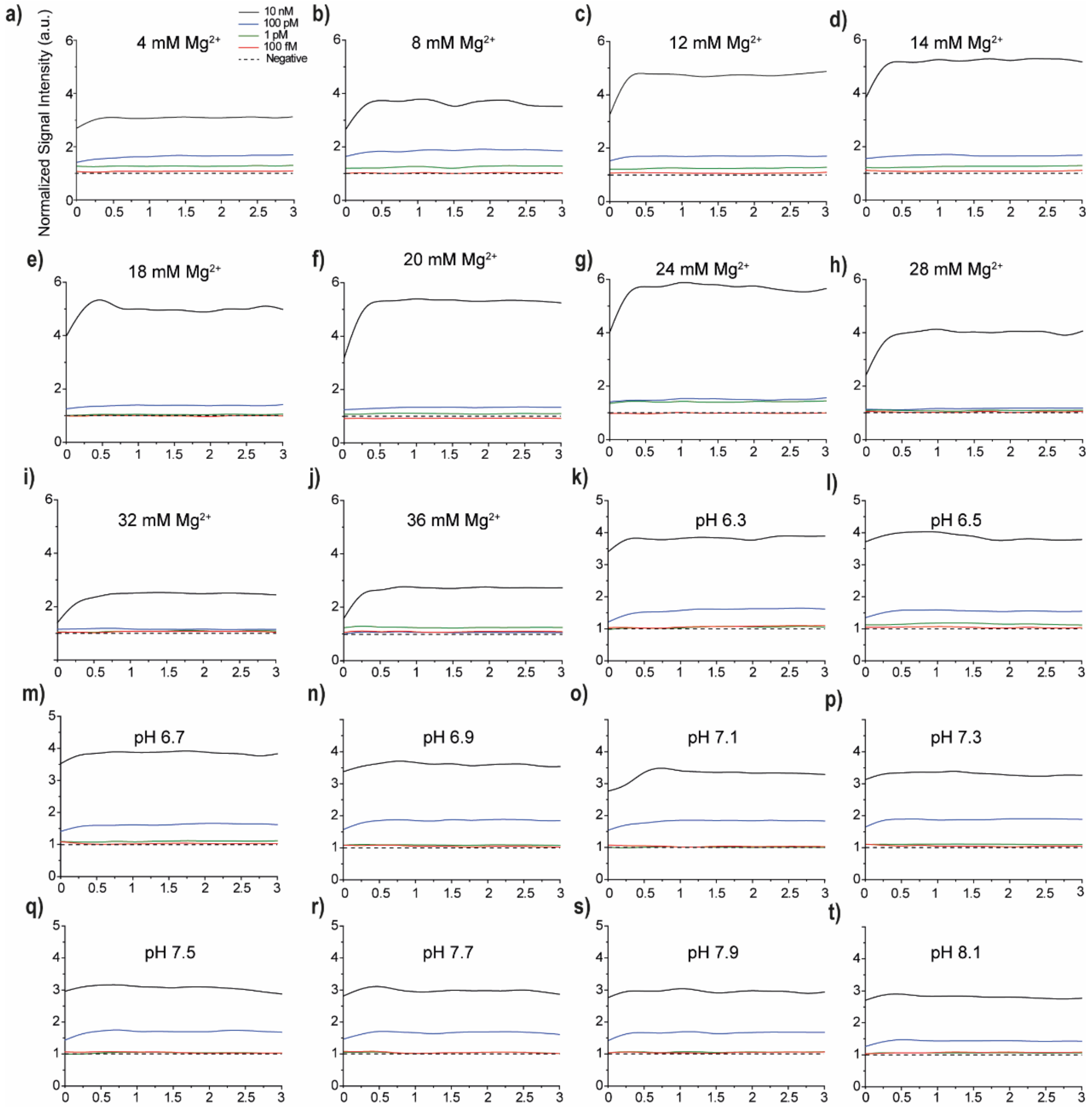

**Supplementary Figure 3. Optimization of CRISPR/Cas13a assay with $Mg^{2+}$ and pH. a-j)** $Mg^{2+}$ concentration from 4 mM to 36 mM. **k-t)** Different pH for off-coverslip assay from pH 6.3 to pH 8.1 by step 0.2.

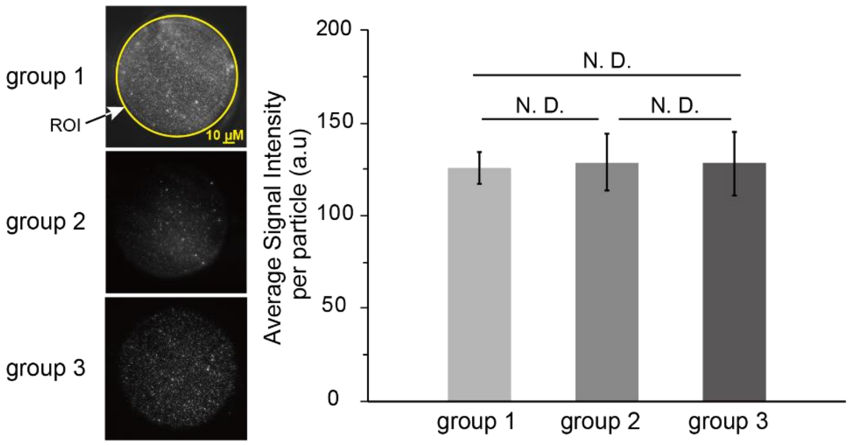


**Supplementary Figure 4.** The average intensity per fluorescent particle from random TIRF images group 1 to group 3.

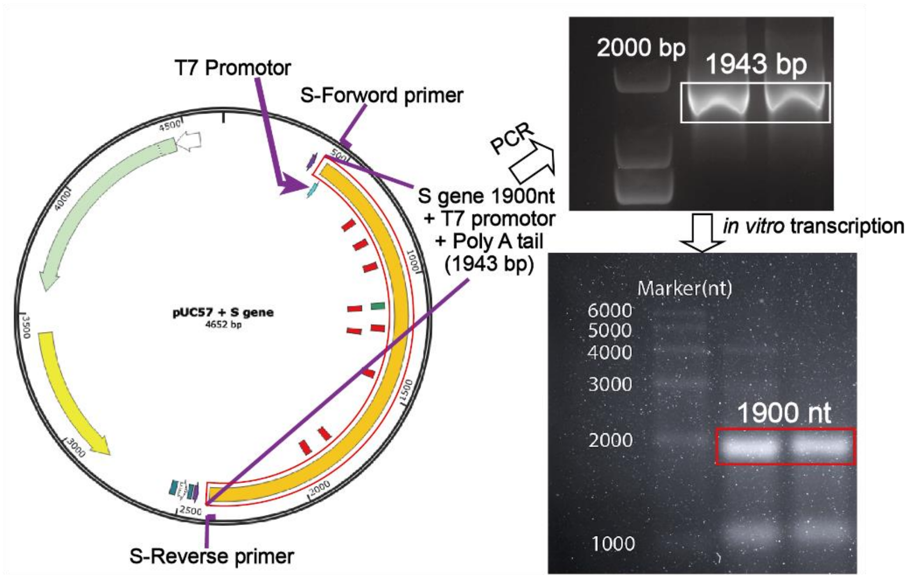


**Supplementary Figure 5**. The plasmid, PCR, and *in vitro* transcription of SARS-CoV-2 S gene. **a)** The pUC57 plasmid contains 1900 nt S gene template, T7 promotor, and polyA tail. **b)** The amplified target DNA fragment. The agarose gel shows the size of the amplified target DNA is about 1900 bp consistent with the theoretical size. **c)** RNA fragment of S gene. The transcription of 1900 nt S gene RNA is confirmed by denaturing formaldehyde agarose gel.

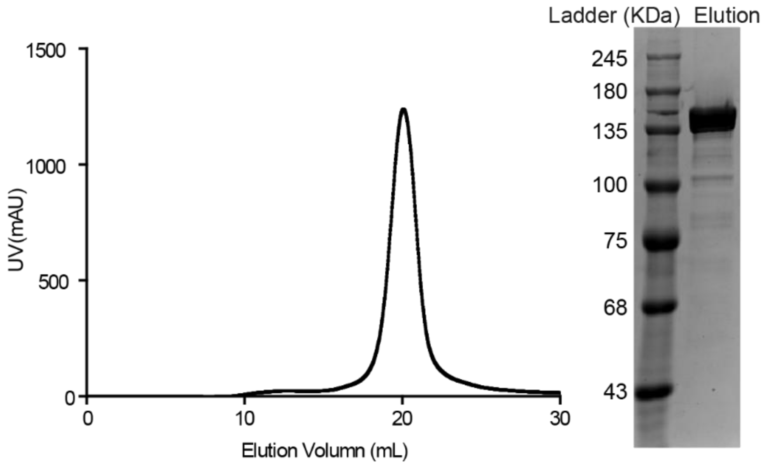


**Supplementary Figure 6**. FPLC chromatograms: a representative chromatogram of cation exchange (Mono S, GE Healthcare Life Sciences) *Lwa*Cas13a. The UV absorbance in milli-arbitrary units (mAU) is plotted against the elution volume in milliliters. Coomassie-stained SDS-PAGE gel is used to confirm the size of *Lwa*Cas13a after elution from cation exchange chromatography.

Supplementary Table 1: crRNA used in this study.

| | |
|---|---|
| 5U reporter | 5’ 6FAM-UUUUU-BHQ1 3’ |
| 8U reporter | 5’ 6FAM-UUUUUUUU-BHQ1 3’ |
| 14U reporter | 5’ 6FAM-UUUUUUUUUUUUUU-BHQ1 3’ |
| 18U reporter | 5’ 6FAM-UUUUUUUUUUUUUUUUUU-BHQ1 3’ |
| 22U reporter | 5’ 6FAM-UUUUUUUUUUUUUUUUUUUUUU-BHQ1 3’ |

Supplementary Table 2: Target sequence.

| | |
|---|---|
| Target-1<br>(1900 nt) | 5’-AAUGUUUGUUUUUCUUGUUUUAUUGCCACUAGUCUCUAGUCAGUG<br>UGUUAAUCUUACAACCAGAACUCAAUUACCCCCUGCAUACACUAAUU<br>CUUUCACACGUGGUGUUUAUUACCCUGACAAAGUUUUCAGAUCCUCA<br>GUUUUACAUUCAACUCAGGACUUGUUCUUACCUUUCUUUUCCAAUGU<br>UACUUGGUUCCAUGCUAUACAUGUCUCUGGGACCAAUGGUACUAAGA<br>GGUUUGAUAACCCUGUCCUACCAUUUAAUGAUGGUGUUUAUUUUGCU<br>UCCACUGAGAAGUCUAACAUAAUAAGAGGCUGGAUUUUUGGUACUAC<br>UUUAGAUUCGAAGACCCAGUCCCUACUUAUUGUUAAUAACGCUACUA<br>AUGUUGUUAUUAAAGUCUGUGAAUUUCAAUUUUGUAAUGAUCCAUUU<br>UUGGGUGUUUAUUACCACAAAAACAACAAAAGUUGGAUGGAAAGUGA<br>GUUCAGAGUUUAUUCUAGUGCGAAUAAUUGCACUUUUGAAUAUGUCU<br>CUCAGCCUUUUCUUAUGGACCUUGAAGGAAAACAGGGUAAUUUCAAA<br>AAUCUUAGGGAAUUUGUGUUUAAGAAUAUUGAUGGUUAUUUUAAAA<br>UAUAUUCUAAGCACACGCCUAUUAAUUUAGUGCGUGAUCUCCCUCAG<br>GGUUUUUCGGCUUUAGAACCAUUGGUAGAUUUGCCAAUAGGUAUUAA<br>CAUCACUAGGUUUCAAACUUUACUUGCUUUACAUAGAAGUUAUUUGA<br>CUCCUGGUGAUUCUUCUUCAGGUUGGACAGCUGGUGCUGCAGCUUAU<br>UAUGUGGGUUAUCUUCAACCUAGGACUUUUCUAUUAAAAUAUAAUGA<br>AAAUGGAACCAUUACAGAUGCUGUAGACUGUGCACUUGACCCUCUCU<br>CAGAAACAAAGUGUACGUUGAAAUCCUUCACUGUAGAAAAAGGAAUC<br>UAUCAAACUUCUAACUUUAGAGUCCAACCAACAGAAUCUAUUGUUAG<br>AUUUCCUAAUAUUACAAACUUGUGCCCUUUUGGUGAAGUUUUUAACG<br>CCACCAGAUUUGCAUCUGUUUAUGCUUGGAACAGGAAGAGAAUCAGC<br>AACUGUGUUGCUGAUUAUUCUGUCCUAUAUAAUUCCGCAUCAUUUUC<br>CACUUUUAAGUGUUAUGGAGUGUCUCCUACUAAAUUAAAUGAUCUCU<br>GCUUUACUAAUGUCUAUGCAGAUUCAUUUGUAAUUAGAGGUGAUGAA<br>GUCAGACAAAUCGCUCCAGGGCAAACUGGAAAGAUUGCUGAUUAUAA<br>UUAUAAAUUACCAGAUGAUUUUACAGGCUGCGUUAUAGCUUGGAAUU<br>CUAACAAUCUUGAUUCUAAGGUUGGUGGUAAUUAUAAUUACCUGUAU<br>AGAUUGUUUAGGAAGUCUAAUCUCAAACCUUUUGAGAGAGAUAUUUC<br>AACUGAAAUCUAUCAGGCCGGUAGCACACCUUGUAAUGGUGUUGAAG<br>GUUUUAAUUGUUACUUUCCUUUACAAUCAUAUGGUUUCCAACCCACU |

| | |
|---|---|
| | AAUGGUGUUGGUUACCAACCAUACAGAGUAGUAGUACUUUCUUUUGA ACUUCUACAUGCACCAGCAACUGUUUGUGGACCUAAAAAGUCUACUA AUUUGGUUAAAAACAAAUGUGUCAAUUUCAACUUCAAUGGUUUAACA GGCACAGGUGUUCUUACUGAGUCUAACAAAAAGUUUCUGCCUUUCCA ACAAUUUGGCAGAGACAUUGCUGACACUACUGAUGCUGUCCGUGAUC CACAGACACUUGAGAUUCUUGACAUUACACCAUGUUCUUUUGGUGGU GUCAGUGUUAUAACACCAGGAACAAAUACUUCUAACCAGGUUGCUGU UCUUUAUCAGGAUGUUAACUGCACAGAAGUCCCUGUUGCUAUUCAUG CAGAUCAACUUA-3’ |
| Target-2 (99 nt) | 5’-AACUUUACUUGCUUUACAUAGAAGUUAUUUGACUCCUGGUGAUUC UUCUUCAGGUUGGACAGCUGGUGCUGCAGCUUAUUAUGUGGGUUAUC UUCAACCU-3’ |

Supplementary Table 3: crRNAs used in this study

| | |
|---|---|
| crRNA-1 | 5’-gauuuagacuaccccaaaaacgaaggggacuaaaac UAACAAUAGAUUCUGUUGGUUGGACUCUA-3’ |
| crRNA-2 | 5’-gauuuagacuaccccaaaaacgaaggggacuaaaac GUAGGGACUGGGUCUUCGAAUCUAAAGUA-3’ |
| crRNA-3 | 5’-gauuuagacuaccccaaaaacgaaggggacuaaaac GCCGAAAAACCCUGAGGGAGAUCACGCAC-3’ |
| crRNA-4 | 5’-gauuuagacuaccccaaaaacgaaggggacuaaaac AAACCCUGAGGGAGAUCACGCACUAAAUU-3’ |
| crRNA-5 | 5’-gauuuagacuaccccaaaaacgaaggggacuaaaac GAAGAAUCACCAGGAGUCAAAUAACUUCU-3’ |
| crRNA-6 | 5’-gauuuagacuaccccaaaaacgaaggggacuaaaac UAUCAAACCUCUUAGUACCAUUGGUCC-3’ |
| crRNA-7 | 5’-gauuuagacuaccccaaaaacgaaggggacuaaaac AACUCACUUUCCAUCCAACUUUUGUUGUU-3’ |
| crRNA-8 | 5’-gauuuagacuaccccaaaaacgaaggggacuaaaac UUACCACCAACCUUAGAAUCAAGAUUGUU-3’ |
| crRNA-9 | 5’-gauuuagacuaccccaaaaacgaaggggacuaaaac AAACCUUCAACACCAUUACAAGGUGUGCU-3’ |
| crRNA-10 | 5’-gauuuagacuaccccaaaaacgaaggggacuaaaac UGCAGCACCAGCUGUCCAACCUGAAG-3’ |

*The lower cases represent the 36-nt repeat region for LwaCas13a and the capital cases represent the 28-nt guide region.
